# Physiological predictors of reproductive performance in the European Starling (*Sturnus vulgaris*): I. Univariate analysis

**DOI:** 10.1101/332577

**Authors:** M.A. Fowler, A. Cohen, M. Paquet, T.D. Williams

**Affiliations:** Simon Fraser University, Department of Biological Sciences, 8888 University Dr., Burnaby, BC, V5A 1S6, Canada (TDW e-mail); Springfield College. Biology, 263 Alden Street, Springfield, MA 01109-3797; Groupe de recherche PRIMUS, Dept. of Family Medicine, University of Sherbrooke, 3001 12e Ave N, Sherbrooke, QC, J1H 5N4, Canada (AAC email)

**Keywords:** European starling, *Sturnus vulgaris*, physiological state, reproductive fitness

## Abstract

Costs of reproduction are assumed to be widespread, and to have a physiological basis, but the mechanisms driving decreased survival and future fecundity remain elusive. Our overall goals were to assess whether physiological variables are related to workload ability or immediate fitness consequences and if they mediate future survival or reproductive success. We investigated individual variation in physiological state at two breeding stages (incubation, chick-rearing) in female European starlings (*Sturnus vulgaris*), for first-and second-broods over two years. Specifically, we measured a suite of 13 physiological traits related to aerobic/metabolic capacity, oxidative stress and muscle damage, intermediary metabolism and energy supply, and immune function. We tested for relationships to traits for workload (e.g. nest visit rate) and fitness (current reproduction, survival and future reproduction).

There was little co-variation among the 13 physiological traits, either in incubating or chick-rearing birds. There were some systematic differences in incubation versus chick-rearing physiology. Chick-rearing birds had lower hematocrit and plasma creatine kinase but higher hemoglobin, triglyceride and uric acid levels. Only plasma corticosterone was repeatable between incubation and chick-rearing. We assessed relationships in a univariate manner, and found very few significant relationships between incubation or chick-rearing physiology and measures of workload, current productivity, future fecundity or survival. Variability in ecological context may complicate the relationship between physiology and behavior. Additionally, individuals may regulate different aspects of their physiology independently, making detection of physiological mediators of cost of reproduction difficult. In a companion paper (Cohen et al. submitted) we explore the utility of multivariate analysis (Principal Components Analysis, Mahalanobis distance) of the same data to account for potentially complex physiological integration.

## Introduction

Raising offspring is widely assumed to be costly. Increased activity and energy expenditure are often associated with rearing offspring ((1-3), but see (4)) and thought to generate negative physiological consequences (5, 6). Incurring these costs may impact survival or future fecundity, known as a cost of reproduction. This concept of cost of reproduction is a central tenet of life history theory, which predicts that parents will face a trade-off between the cost incurred for the current reproductive event and the potential for future reproductive opportunities (7-9). In order to preserve future reproductive potential and thus maximize lifetime reproductive success, investment in each reproductive event should reflect a physiological optimum, rather than a physiological maximum (2, 10, 11).

Cost of reproduction research is of interest to theoretical ecologists studying life histories, but also has broader applications, as we seek to understand the limitations of organisms and their reproductive patterns. Interactions with humans can impact different aspects of physiology and parental care (12-15), which if costly, has the potential to affect long term reproductive patterns and demographics of populations (11). Therefore, gathering data quantifying these costs of reproduction has value on multiple scales. Despite the importance of understanding the mechanisms behind costs of reproduction, questions remain about what actually drives these costs (7, 16, 17).

There is considerable variation in cost of reproduction in the avian literature. Parental workload (offspring provisioning) varies widely (18, 19), but the question of what mechanism(s) determines individual variation in workload remains very poorly resolved (20, 21). The physiological consequences of egg production have been documented (22) and there is potential for physiological costs due to offspring provisioning (5, 6, 23). Many studies have focused on energy expenditure, widely considered to be the currency of life-histories even though this is only one component of the complex physiology of free-living animals, but these have failed to reveal clear relationships with variation in parental care (24). Some correlational studies have shown that variation in single physiological measures can be systematically related to workload or aerobic capacity ((25-27), but see (28)). Other studies have measured multiple physiological traits (29, 30), including in experimental studies (31). However, these studies reveal few, or complex relationships between workload (e.g. nestling feeding frequency) and physiology.

The equivocal results in avian systems indicate that physiological costs of reproduction may not be manifest in a single breeding attempt. Additionally, there may be little covariation among physiological systems (e.g. immune function, Matson et al. (32) or antioxidant levels, Cohen and McGraw (33)). Birds may be able to adjust individual components of their physiology independently (34-36) or there may be complex, context-dependent relationships among components (37). Data from multiple breeding attempts and a suite of physiological variables from multiple systems may shed light on these complex relationships.

In this paper we analyse individual variation in physiological state in relation to parental care at two breeding stages (incubation and chick-rearing) in female European starlings (*Sturnus vulgaris*), for first-and second-broods in two years, using a repeated-measures design. European starlings are an ideal system in that they readily use nest boxes at our field site and are highly synchronous breeders, minimizing variability due to time of year (20). We measured 13 physiological variables at both stages, as well as quantified parental workload (nest visit rate), reproductive success and survival. We assess the predictive capacity of these physiological variables on two categories of response variables: workload and fitness costs (immediate or future). Workload variables reflect how hard females were working: nest visit rate and brood size she must provision during chick-rearing. The fitness costs indicate potential life history consequences: current reproduction, future fecundity (in second brood or in second year), survival and cumulative productivity (over 2 years) (Table 1).

**Table 1.** Summary of physiological and morphological predictor variables and life-history outcome variables.

| **Physiological/morphological traits** | **Outcome variables** |
| --- | --- |
| **<u>Aerobic/metabolic capacity</u>** | **<u>Workload</u>** |
| Hematocrit (Hct) | Nest visit (provisioning) rate |
| Hemoglobin (Hb) | Brood size at day 6 (BS6) |
| Reticulocytes |  |
| Corticosterone (Cort) |  |
| **<u>Oxidative stress/muscle damage</u>** | **<u>Current reproduction</u>** |
| Reactive oxygen metabolites (dROMs) | Total failure/success (incubation only) |
| Total antioxidants (OXY) | Brood size at fledging, brood 1 |
| Creatine kinase (CK) | Mean chick fledging mass, brood 1 |
|  | Year 1 total offspring |
| **<u>Intermediary metabolism/energy supply</u>** | **<u>Future fecundity</u>** |
| Non-esterified fatty acids (NEFA) | % initiating 2<sup>nd</sup> brood |
| Triglycerides | Brood size at fledging, brood 2 |
| Uric acid | Mean fledging mass, brood 2 |
| Glucose | BSF for brood 1 in year 2 |
| **<u>Immune function</u>** | **<u>Survival</u>** |
| Haptoglobin | Local return rate in year 2 |
| Natural antibodies (NAb) |  |
| **<u>Morphological traits</u>** | **<u>Cumulative productivity</u>** |
| Body mass | Cumulative (2 year) BSF |
| Year-centered wing loading |  |

Studies have suggested that avian costs of reproduction may be associated with aerobic capacity (21), oxidative stress (38), fuel use (30), and immune function (39). Thus, the physiological variables we chose reflect four general physiological components: 1) aerobic/metabolic capacity, 2) oxidative stress and muscle damage, 3) intermediary metabolism and energy supply, and 4) immune function (Table 1). Aerobic capacity/metabolic capacity was encompassed by the following physiological variables: hematocrit (Hct), hemoglobin (Hb), reticulocytes, and corticosterone (cort). Oxidative stress and muscle damage variables included: reactive oxygen metabolites (dROMs), total antioxidants (OXY) and creatine kinase (CK). Intermediary metabolism and energy supply was assessed by quantifying: non-esterified fatty acids (NEFA), triglycerides, uric acid and glucose. Immune function was reflected by the measurement of: haptoglobin and natural antibodies (Table 1).

Our overall goals were to assess whether the physiological variables are related to work load ability or immediate fitness consequences and if they mediate future survival or reproductive success. Within this broad framework, we had several specific aims:

1. First, we consider the correlations among the suite of physiological traits in both incubating and chick-rearing females, the change of these traits between stages and repeatability.
2. We then test whether physiological state during chick-rearing is predictive of the workload for the current brood or is related to current or future fitness metrics.
3. Finally, we incorporate not only the absolute physiological values, but also the change (delta) in physiological variables from incubation to chick-rearing, potentially reflecting physiological adjustments individuals make. We investigate the power of delta incubation:chick-rearing physiology to predict workload or fitness costs (immediate or future).

Several analytical techniques can be applied to these data. In this paper, we present univariate analyses in order to test biologically intuitive hypotheses, as well as facilitate interpretation of the relationships. However, there is a growing recognition of the complex and integrative nature of physiological systems (37, 40, 41). In a companion paper (Cohen et al, submitted) we explore the utility of analysing the same data set using multivariate analysis (Principal Components Analysis, Mahalanobis distance) to account for potentially complex physiological integration.

## Methods

Fieldwork was conducted at Davistead Farm, Langley, British Columbia, Canada (49°10′N, 122°50′W) between April-early July 2013-2014 and females were followed in 2014 and 2015 to measure maternal survival (local return rate) and fecundity in the year after blood sampling. This site comprises about 150 nest boxes mounted on posts around pastures and on farm buildings. Nest boxes were checked daily from April 1 to determine laying date, clutch size, brood size at hatching, at day 6 post-hatching, and at fledging (day 21), and mass and tarsus of all chicks was recorded when chicks were 17 days old. We then obtained the same data for second broods.

All blood samples were collected by puncturing the brachial vein with a 26½-gauge needle and collecting blood (<700 μL) into heparinized capillary tubes. At the same time fresh blood was collected for a) hematocrit and hemoglobin measurements, b) two blood smears were prepared for reticulocyte counts, and c) glucose levels (mmol.L^−1^) were measured with a handheld glucose meter (Accu-chek Aviva®; see Supplemental Material S1). Blood samples were stored at 4°C for up to 4 h before being centrifuged for 6 min at 10,000*g*, plasma was collected and then stored at −80°C until assayed. All research was conducted under Simon Fraser University Animal Care permits # 657B-96, 829B-96, 1018B-96).

Although females were captured using different methods in incubation and chick-rearing (see below) all birds were blood sampled with 3 minutes of being handled. Incubation (day 6-8) samples from all breeding females were obtained by plugging the nest box hole prior to dawn. This method can result in different time periods that the birds are passively sitting in the nest box (mean 56 minutes, maximum 132 minutes) before being removed and blood sampled. However, there was no relationship between time in box and plasma corticosterone levels, oxidative stress, or immune function, suggesting that birds did not perceive this as a stressor (Cohen et al, submitted). Incubating females were fitted with color bands and individually numbered metal bands (Environment Canada permit # 10646; males were not captured or banded, and thus, identity of males is unknown). As many females as possible were recaptured during chick-rearing of first broods (day 10-12) and again during chick-rearing of second broods (day 10-12). For a sub-sample of females (2013, *n* = 12; 2014, *n* = 14) we attached radio transmitters (Holohil Limited, Inc., model BD-2, mass = 1.9g; data included in the companion paper). There was no difference in mean brood size at fledgingmean 17-day chick mass, mean provisioning rate, or any physiological trait for females with and without radio-transmitters (*P* > 0.05 in all cases) so we pooled data for all subsequent analyses. During chick-rearing females were caught using nest traps (Van Ert Enterprises, Leon, IA) as they entered nest boxes to feed chicks, removed immediately and blood sampled within 3 minutes.

### Measurement of physiological traits

We analysed plasma samples for 13 physiological traits using standard assay methods reported in previous studies; see Supplemental Material S1 for full details. Due to variation in the amount of plasma we obtained, for some individuals there was not enough plasma to run all assays, thus sample sizes differ for some physiological traits. For all traits except Natural antibodies (NAb) we used either raw data values or square-root or natural log transformed data (see Supplemental Material S1 and Supplemental Tables). For Natural antibody (NAb) immune measures agglutination and lysis scores were positively correlated and we chose to combine them with principal component analysis, thus reducing the total variables. This variable is referred to in the analysis as NAb PC1.

### Wing area measurements and wing loading

Wing surface area (chick rearing only) was calculated from digital photos taken in the field using the free image software IMAGEJ (available from http://rsbweb.nih.gov/ij/). We present chick-rearing wingloading only, as we were interested in the potential flight effects of provisioning, but were less interested in the flight ability of birds incubating eggs. Two wing photos were taken of the left wing spread on a board with 2 cm grid drawn. Each picture was scaled to the 2 cm grid using the software. The outline of the wing was traced in IMAGEJ two times for each photo, resulting in four measurements per individual. The coefficient of variation between measures in one photo was 0.39% and between two photos was 3.13%. Wing surface areas were averaged and doubled to attain total wing surface area. The body box (Pennycuick 2008, i.e the area between the wings) was not included in the calculation of the wing surface area. Wing loading was calculated as mass/area (42). Due to a difference in measurers by year, the control, chick-rearing values are mean centered within year.

### Measures of workload and fitness costs

We used two measures of workload, reflecting how hard the female must work (Table 1). Nest visit rate or provisioning rate and day 6 brood size (i.e. brood demand during the linear phase of chick growth (43)). Nest visit rate was obtained from 30-minute behavioral observations surveys between 09.00-14.00 on days 6, 7, and 8 post-hatching with 2-3 observations per nest (i.e. either 1-hour or 1.5-hours of data per nest; Fowler & Williams 2015). Days 6-8 were chosen as they represent the period of most rapid chick growth (43) and represents more chicks that require nutrition. We recorded number of visits for males and females separately, based on presence of bands/colour bands. Visits where sex could not be determined were initially categorized as unknown, but unknown visits were quite low (∼5-15% of total observations). Unknown visits were then partitioned between male and females based on the ratio of known visits, after Fowler and Williams (18).

Fitness metrics included current and future reproductive success, as well as survival. Specifically, the variables we used included current reproduction as a) brood size at fledging (day 21 post-hatching) and b) mean chick fledging mass (measured at day 17), for the first brood. In addition, for incubating birds only, we compared physiological traits among birds with total breeding failure or breeding success (≥ 1 chick fledged), and brood size at day 6 post-hatching, to assess immediate consequences of variation in incubation physiology. As measures of future fecundity we used, a) probability of initiating a second clutch (0/1), b) brood size at fledging (day 21 post-hatching) and mean chick fledging mass, for the second brood, and c) brood size at fledging for the brood in year 2. We present the percent of individuals initiating 2^nd^ broods, but model the data as a logistic regression. Finally, we compared physiological traits during chick-rearing in year 1 to local return rate in year 2, and to the cumulative number of chicks fledged over 2 years in all breeding attempts (see Table 1).

### Statistical analysis

All data were analyzed using R version 3.2.1 (R Core Team 2015). We tested for normality and normalized all non-normal variables using either natural log or square root transformation (see Supplemental Material S1). After normalizing data we tested for, but did not detect, any statistical outliers for any physiological traits except Cort so we did not exclude any data for other physiological traits. For plasma Cort where we excluded n = 7 females which had Cort > 80 ng/ml, which is within the range of “stress-induced” levels in this species (44). These samples showed no relationship to time in box. For individuals where we had physiological trait values during incubation and chick-rearing (for the first brood) we calculated the change in trait value (delta or Δ) between stages. For analyses of correlations among multiple traits, in incubating and chick-rearing birds, we considered results both using raw P values (P < 0.05) for exploratory purposes, and using adjusted P values in R based on False Discovery Rate using the Benjamini/Hochberg correction (45).

Repeatability estimates were generated with the rptR package (46) and individual included as random after (18). Physiological changes between incubation and chick-rearing were assessed with linear mixed effects models, including band and year as random effects. Analyses of physiological predictors of workload and cost of workload were conducted using lme4 package (47) with year and individual identity as random effects and body mass as covariate. P values were calculated with the Kenward-Rogers correction (48). When brood size was investigated as a response variable, we used generalized mixed effects models with Poisson error distributions, and report the z-statistic and associated P value. Binomial response variables (success, 2^nd^ brood initiation, or survival) were analyzed as logistic regression.

## Results

Our maximum samples sizes of female European starlings were as follows: a) incubation, 2013 *n* = 46 and 2014; *n* = 30, total *n* = 76; b) chick-rearing, first brood, 2013 *n* = 31 and 2014 *n* = 24, total *n* = 55; and c) chick-rearing second brood, 2013 *n* = 14 and 2014 *n* = 7, total *n* = 21. Here, we restrict our analysis of physiological data to samples obtained during incubation and chick-rearing for first broods due to the relatively small number of second brood samples (but see companion paper, Cohen et al. submitted). Raw data values (mean ± S.D.) for all physiological traits by year and breeding stage are given in Supplemental Table S1.

### Aim 1: Correlations among the suite of physiological traits in both incubating and chick-rearing females, change between stages and repeatability

#### Covariation among physiological traits in incubating females

Only 3/13 traits were significantly correlated with body mass: number of reticulocytes (*r*70 = - 0.25, *P* = 0.040) and plasma total antioxidants (*r*73 = −0.26, *P* = 0.024) were negatively correlated with mass, and plasma haptoglobin was positively correlated (*r*70 = 0.31, *P* = 0.009) but all traits were independent of body mass at the FDR corrected *P* value (Supplemental Table S2).

Among 13 physiological traits, 21 pairwise correlations were significant at *P* < 0.05 (Supplemental Table S2). Within functional groups (at *P* < 0.05), a) hematocrit and hemoglobin were positively correlated, b) dROMs and OXY were strongly positive correlated and both were negatively correlated with plasma CK and haptoglobin; c) NEFA, triglyceride and uric acid were positively correlated, but plasma glucose was only positively correlated with Uric acid; and d) haptoglobin and NAb were negatively correlated (Supplemental Table S1). Haptoglobin was correlated with 7/13 traits in incubating females (at *P* < 0.05): negatively with hemoglobin, reticulocytes, dROMs and OXY, and positively with creatine kinase, NEFA and uric acid. Using the FDR corrected *P* value only 6 pairwise correlations among physiological traits were significant. dROMs correlated positively with OXY and negatively with uric acid, and haptoglobin correlated negatively with reticulocytes, dROMs, OXY, and positively with CK (all *P* < 0.01).

#### Covariation among physiological traits in chick-rearing females

All 13 physiological traits were independent of body mass (*P* > 0.05 in all cases) and year-centered wing loading (*P* > 0.05 in all cases; Supplemental Table S3). Ten pairwise correlations were significant at *P* < 0.05. Similar to incubation, (at P < 0.05) hematocrit and hemoglobin were positively correlated (*r*54 = 0.42), as well as dROMs and Oxy (*r*54 = 0.36). Both Oxy (*r*_*52*_ = −0.33) and dROMS (*r*_*51*_ = −0.53) were negatively correlated with haptoglobin. However, there were fewer significant relationships among intermediary metabolites in chick-rearing birds (Supplemental Table S3): plasma glucose was negatively correlated with uric acid (*r*_*54*_ = −0.32), and NEFA (*r*_*54*_ = −0.32). At the FDR corrected P value (which might be overly conservative), only two relationships remained significant: hematocrit and hemoglobin (*r*_54_ = 0.42), and haptoglobin and dROMs (*r*_51_= −0.53).

#### Changes between Incubation and Chick-rearing

For three physiological traits there was a year*breeding stage interaction (*P* < 0.001).

Reticulocyte counts decreased from 15.6 ± 6.1% in incubating birds to 8.2 ± 5.1% during chick-rearing in 2013 but did not change between incubation (8.7 ± 5.9%) and chick-rearing (7.3 ± 2.5%) in 2014 (Fig. 1a). Secondly, blood glucose decreased between breeding stages in 2013 and increased between stages in 2014 (Fig. 1b). Finally, OXY increased from incubation to chick-rearing in both years but the change was much greater in 2014 (167.6 vs 232.9 μmol HClO.ml^−1^; +39.0% compared with 2013 (254.4 vs. 271.1μmol HClO.ml^−1^; +6.6%).

**Figure 1.**
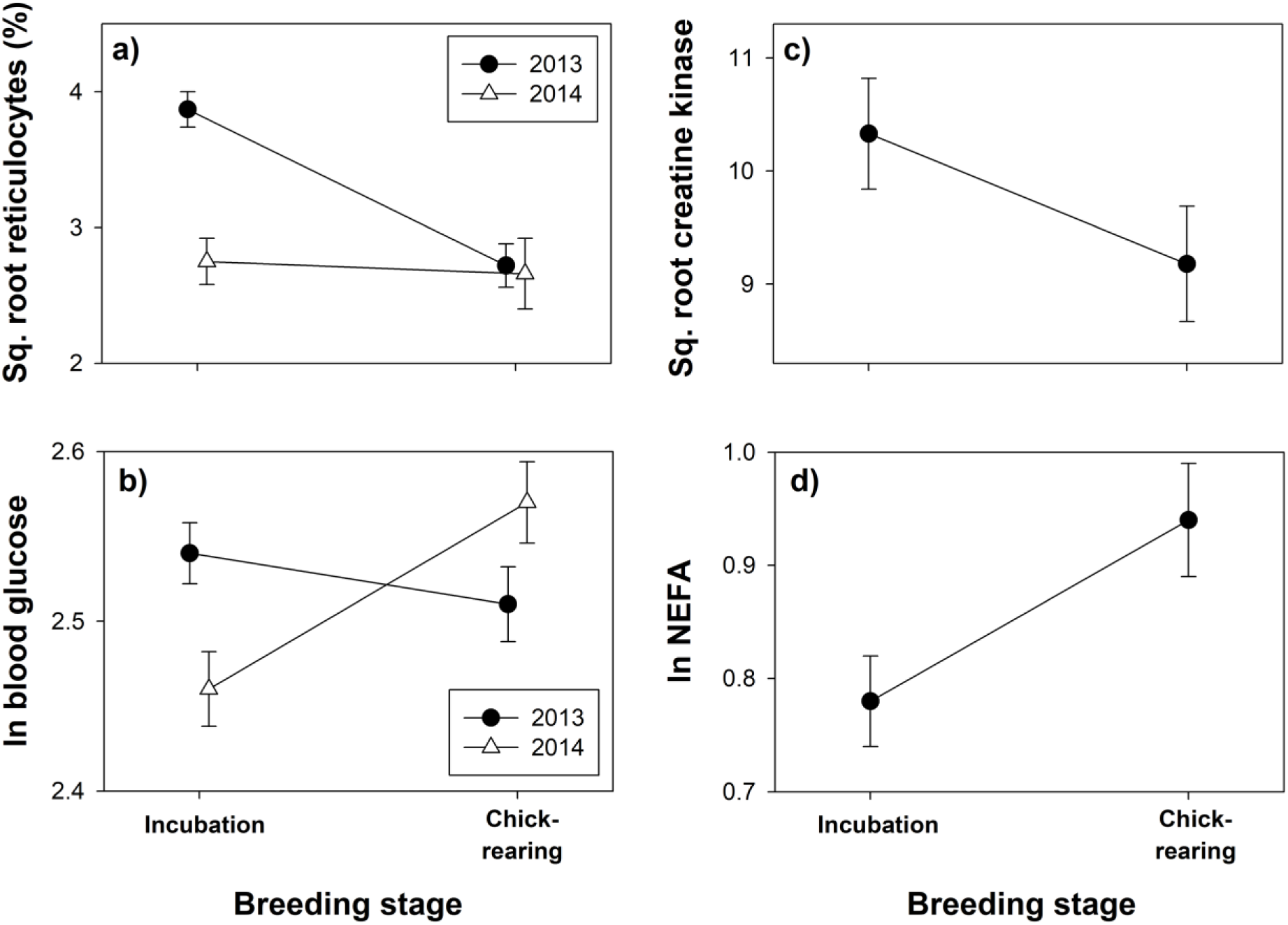
Representative examples of patterns of change in physiological traits showing breeding stage*year interactions: a) reticulocyte counts, and b) blood glucose levels, and traits showing an effect of breeding stage with no interaction: c) plasma creatine kinase and d) plasma non-esterified fatty acids; see text for details.

For six traits there was a significant effect of year, but no year*breeding stage interaction (P > 0.05): hemoglobin (P < 0.01), CK (p = 0.04), dROMs (P < 0.001), NEFA (p = 0.04), uric acid (P = 0.03) and haptoglobin (P < 0.001). We therefore included “year” as a random factor in analysis of breeding stage differences for the 10 traits with no year*stage interaction. Chick-rearing birds had lower body mass (P = 0.04), hematocrit (P = 0.03), dROMs (P = 0.03), and plasma CK (P < 0.01; Fig. 1c) but higher hemoglobin (P = 0.03), NEFA (Fig. 1d), triglyceride and uric acid (all P < 0.001). There was no difference in log Cort, haptoglobin or NAb PC1 (all P > 0.05) among breeding stages or years.

The only physiological trait that was repeatable between incubation and chick-rearing in *both* years was plasma corticosterone (2013, *r* = 0.67, *P* = 0.03; 2014, *r* = 0.69, *P* < 0.001; overall *r* = 0.55; Fig. 2a). Hematocrit was repeatable in 2013 (*r* = 0.42, *P* = 0.01) but not in 2014 (*r* = 0.03, *P* = 0.09; Fig. 2b). Similarly, creatine kinase was highly repeatable in 2013 (*r* = 0.75, *P* < 0.001) but not in 2014 (*r* = 0, *P* > 0.70; Fig. 2c). For some traits year differences would have generated ‘apparent’ repeatability if pooled data were analysed (e.g. Fig. 2d).

**Figure 2.**
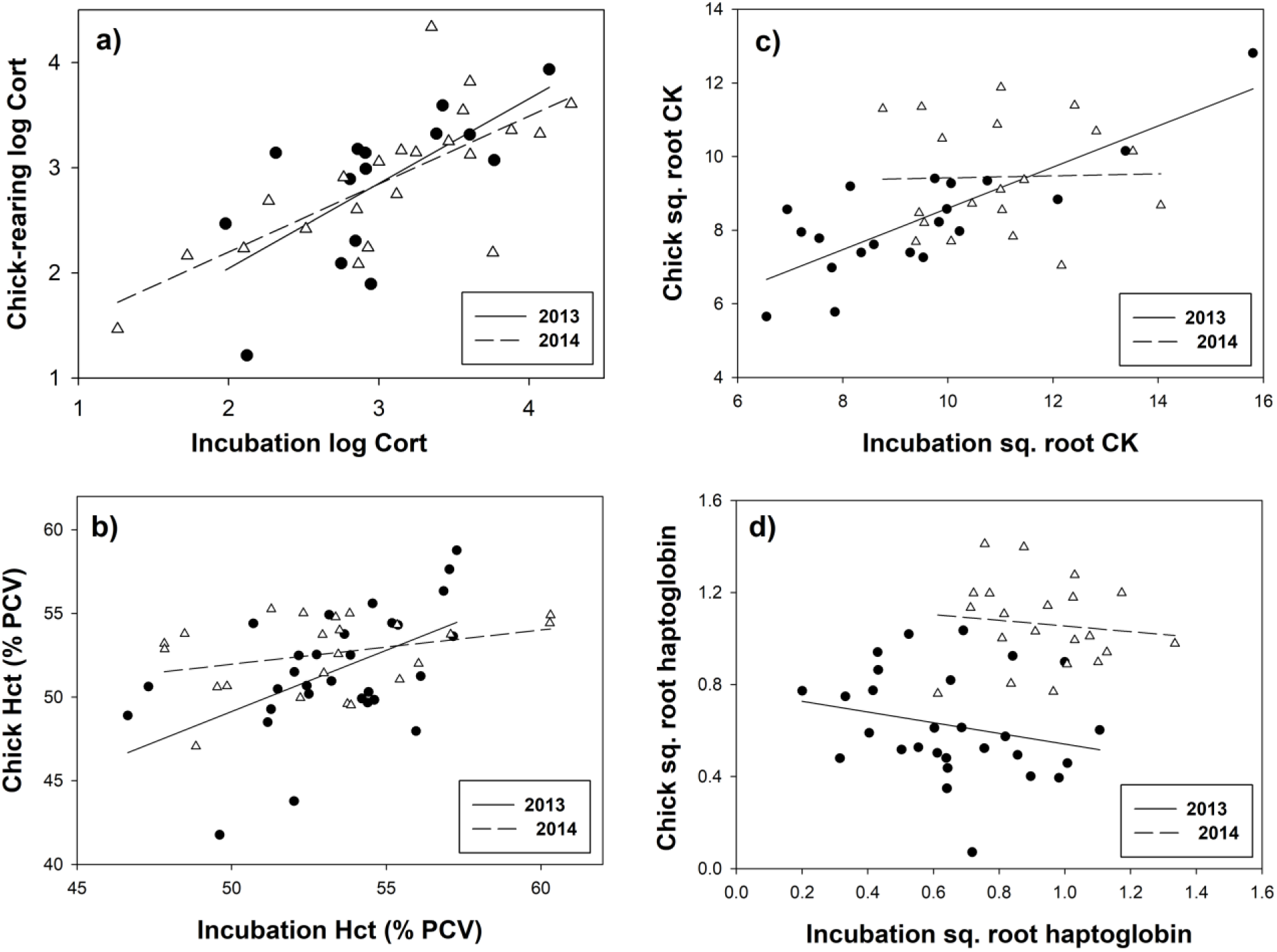
Representative examples of patterns of repeatability in physiological traits: a) plasma corticosterone, b) hematocrit, c) plasma creatine kinase, and d) haptoglobin; see text for details.

### Aim 2) Relationship between physiological state and workload and fitness costs

We found that physiological state during incubation did not predict BS6 (P > 0.05 in all cases), demonstrating that incubation physiology does not reflect immediate consequences. Additionally, there was no difference in physiological trait values for incubating birds among individuals that subsequently showed total breeding failure versus breeding success (≥ 1 chick; logistic regression, P > 0.14 in all cases). This analysis further illustrates that differences in physiology among breeding stages were not related to the fact that some birds were sampled during incubation but were not re-sampled during chick-rearing.

There were significant relationships for only 5/156 (3.2%) pair-wise contrasts between chick-rearing physiological traits and life-history outcomes (at P < 0.05). (see Table 2 for summary of results). For measures of current workload, nest visit (provisioning) rate was positively associated only with plasma corticosterone (*F*_1,9_ = 7841, *n* = 43, *P*< 0.001. For measures of current reproduction mean fledging mass for the current brood (brood 1) was negatively associated with NEFA (F1,46.1 = 6.6, n = 50, P = 0.01). There was a negative relationship between dROMs and the number of chicks fledged from brood 1 (z = −1.95, n = 53, P = 0.047) as well as the cumulative number of chicks fledged from brood 1 and 2 in the current breeding attempt (z = −2.17, n = 52, P = 0.03). Similarly, cumulative BSF was positively associated with triglyceride levels (z = 2.1, p = 0.04). However, none of these relationships between productivity and either dROMS or triglyceride were significant (P > 0.05) if the three females with BSF = 0 were excluded from the analysis. Finally, there were no significant relationships between any physiological traits and any measure of future fecundity, survival and cumulative productivity (P ≥0.05 in all cases; Table 2).

**Table 2.** Summary of relationships between chick-rearing physiology, workload, current productivity, future fecundity and survival.

| Life-history stage | Outcome variable | Absolute physiological trait value (chick-rearing) | Change in physiological trait value (incubation-chick rearing) |
| --- | --- | --- | --- |
| Workload | Nest visit rate | Cort (+) | Mass (-) |
|  | Brood size at day 6 | - | - |
| Current productivity | Brood size at fledging, brood 1 | dROM (-) <sup>1</sup> | - |
|  | Mean fledging mass, brood 1 | NEFA (-) | Trig (-) |
|  | Year 1 total offspring | dROM (-) <sup>1</sup> | Trig (+) <sup>1</sup> |
| Future fecundity | Probability initiating 2<sup>nd</sup> brood | - | - |
|  | Brood size at fledging, brood 2 | - | OXY (+) <sup>1</sup> |
|  | Mean fledging mass, brood 2 | - | - |
|  | Brood size at fledging for brood 1 in year 2 | - | - |
| Survival | Local return rate, year 2 | - | - |
| Cumulative productivity | No. fledglings over 2 years | Trig (+) <sup>1</sup> | Trig (+) <sup>1</sup> |

<sup>1</sup> Not significant (P > 0.05) when n = 3 females with BSF = 0 are excluded.

### Aim 3) Relationship between delta incubation:chick-rearing and workload or fitness costs

There were significant relationships for only 5/168 (3.0%) pair-wise contrasts of change in physiological trait value and life-history measures (at P < 0.05) (mass was included as a predictive variable rather than a covariate in the delta analysis). Provisioning rate was negatively associated with change in mass between stages (F_1,12.4_ = 6.5, p = 0.03), however individual variation in the change in physiological trait values between incubation and chick-rearing was independent of measures of current workload for all other traits (P ≥ 0.05). The change in triglyceride levels from incubation to chick-rearing was related to immediate fitness consequences in current reproduction. Increasing plasma triglyceride levels were negatively associated with mean fledging mass of chicks in 1st broods (F _1,45.1_ ^=^ 5.9, n = 48, P = 0.02), but was positively related to the cumulative number of chicks fledged from brood 1 and 2 in the current breeding attempt (z = 2.1, n = 52, P = 0.04). Future fitness consequences were also predicted by the change in plasma triglyceride. Increasing triglyceride was also positively related to the cumulative number of chicks fledged over 2 years (z = 2.5, P = 0.01) although sample size here was small (n = 12). Increasing plasma total antioxidant levels (OXY) during the first brood was positively correlated with number of chicks fledged from 2^nd^ broods (z = 2.1, P = 0.04, n = 16). However, once again most of these relationships were not significant (P > 0.05) if the three females with BSF = 0 were excluded from the analysis (Table 2).

## Discussion

Our overall goals were to assess whether physiological variables are related to work load ability or immediate fitness consequences and if they may mediate future fitness costs (survival or reproductive success). To this end, we measured 13 physiological traits, representing four ‘functional groups’ of traits (aerobic/metabolic capacity, oxidative stress and muscle damage, intermediary metabolism and energy supply, and immune function) in both incubating and chick-rearing female European starlings. Surprisingly, we found little evidence for covariation (positive or negative) among our suite of 13 physiological traits either in incubating or chick-rearing birds and even within functional groups of traits, e.g. among intermediary metabolites. Furthermore, physiological traits were largely independent of individual variation in body mass or wing loading. We did find one of the few correlations among a small suite of traits in incubating birds: haptoglobin was negatively correlated with reticulocytes, dROMs and OXY and positively correlated with creatine kinase. Haptoglobin is an acute phase protein that scavenges hemoglobin released into the circulation by hemolysis or normal red blood cell turnover (49, 50). Incubating females are reproductively anemic immediately post-laying (51-53) and this, combined with regenerative erythropoiesis, involves increased red blood cell turnover. Therefore, our results suggest that the effects of reproductive anemia, perhaps mediated by haptoglobin, might extend to several other physiological systems in post-laying females.

We identified systematic differences in incubation physiological metrics relative to chick-rearing individuals. Chick-rearing birds had lower hematocrit and plasma creatine kinase but higher hemoglobin, triglyceride and uric acid levels. However, some of these breeding stage differences varied with year and for three traits (reticulocytes, glucose, total antioxidants) there was a year*breeding stage interaction. Only one trait, plasma corticosterone, was repeatable between incubation and chick-rearing, although hematocrit and creatine kinase were repeatable between breeding stages in 1 of 2 years.

In addition to few correlative relationships among physiological variables, we could predict very little in workload or fitness costs with physiological metrics, either as raw metrics or as the change in physiology between breeding stages. After Williams (54), we investigated whether changes, or physiological reaction norms might be informative. We tested the hypothesis that change in physiological traits from incubation to chick-rearing might be more informative in predicting subsequent performance or reproductive success. Our rationale was that if physiological variation reflected costs of increased activity associated with chick-rearing, then incubation physiology might reflect a base-line, or at least a less active state. Our results do not support these ideas: individual variation in the change in physiology between incubation and chick-rearing was not associated with variation in current or future reproduction or survival. We suggest that this was because birds exhibited a specific incubation physiology, at least for some traits: six of 13 physiological traits were significantly different between breeding stages including hematocrit, hemoglobin, creatine kinase, NEFA, triglyceride and uric acid. Interestingly, our data on variation with breeding stage differ from those of Kern et al. (30) who reported that in female pied flycatchers plasma triglyceride and glucose levels were not different between incubating and chick-rearing birds, but that incubating birds had higher uric acid and free-fatty acid levels. In contrast, we found that chick-rearing birds had higher plasma triglyceride and uric acid levels. Our results further highlight the difficulty of obtaining true baseline physiological values in free-ranging individuals. These individuals were likely experiencing annual variation or demonstrating individual differences in physiological flexibility in different ecological contexts.

We found that with multiple metrics of both physiology and fitness metrics, marked individual variation in life-history traits (reproduction, survival) were independent of large-scale variation in physiological phenotype. This finding is consistent with analyses in our companion paper (Cohen et al.submitted) which suggests minimal efficacy of Principal Component Analysis (PCA) at reducing the dimensionality of our multiple variables due to instability of the correlation structure, and supporting our rationale for exploring variation at the level of univariate traits. Our results, in free-living birds, extends the conclusions from studies by Tieleman et al. (35), Buehler et al. (34) and Versteegh et al. (36) which were based on a smaller number of traits, specific physiological systems, or studies of captive birds. Buehler et al. (34) found no consistent correlations among hematocrit, baseline corticosterone concentration, immune function and basal metabolic rate either within or among individual captive red knot (*Calidris canutus islandica*) subjected to experimentally manipulated temperature treatments over an entire annual cycle. Similarly, Versteegh et al. (36) found considerable phenotypic flexibility among various measures of constitutive immunity (hemagglutination, hemolysis, haptoglobin, bactericidal ability), although four of the five measures of immunity had a genetic component based on a common garden experiment.

Together, these studies suggest that there is little stable covariation not only among physiological components of the same physiological system (e.g. immune function), but also among different physiological systems. The flexibility among physiological components may enable the individual to adjust to a changing environment. Buehler et al. (34) suggested that such physiological flexibility made perfect sense “if one places the animal firmly within its environment where constraints might differ under different circumstances” (see also, (37)). In this sense, the lack of stable covariation in traits does not necessarily indicate a lack of functional relationships, but rather relationships that are highly contingent on individual circumstances. Thus, it is critical to consider both the physiological constraints as well as the ecological context when examining patterns of covariance among phenotypic traits. Versteegh et al. (36) also stressed the importance of considering seasonal variation in immune function in relation to the ecology and life history of the organism of interest (though both these studies were conducted with captive birds).

Our study of free-living birds confirms and highlights the importance of knowing ecological context in studies investigating variation in physiological traits. We found annual variation in physiological trait values, even within breeding stages, lack of repeatability for most traits, and even breeding stage*year interactions. Although we acknowledge that our study was correlational, the two years of this study likely did differ markedly in ecological context. In 2013 mean laying date was relatively early and overall breeding productivity was below-average (a “poor” year), and in 2014 mean laying date was relatively late and breeding productivity was above average (a “good” year) compared with our long-term (15 year means). A growing body of work emphasizes the complexity of the relationship among physiological and performance traits and ecological and life history context (37, 55, 56). Our negative findings don’t reflect only year-to-year variation, they imply that individual variability combined with ecological context likely plays a large role in the interplay between physiological and life history variables.

Physiology as a level of organization has long been a key component of the organismal performance paradigm (57). Careau and Garland (58) designate physiology and biochemistry as lower-level traits and suggest that it is unlikely that these directly influence Darwinian fitness without intermediate effects on performance, behavior, and/or energetics. Indeed, it seems widely accepted that natural selection generally acts most directly on behavior, less on performance abilities, and least directly on lower-level morphological, physiological, and biochemical traits (57-59). The concept that individual variation in physiological traits is filtered through behavior and performance may explain why we were unable to detect direct relationships between physiological traits and life history variables.

There are, however, numerous examples of specific physiological adaptations associated with extreme performance (reviewed in (60). Detecting these associations may depend on whether or not the individuals are motivated to perform maximally and the ecological context in which they must perform a relevant task (61, 62). Despite the correlational nature of our study, we had a very large range of individual variation in the ‘task’ performed, i.e. rearing offspring: some individuals made only one breeding attempt, had no breeding productivity, and did not return the following year, whereas other individuals reared up to 10 chicks from two broods, returned the following year, and reared up to 8 more chicks from two further broods. Accompanying the variation in success, we observed a nine-fold variation in workload (nest visit rate). It is therefore surprising that we see no physiological signature of this marked individual variation in performance and reproductive success. Physiological profiles during incubation did not predict which birds subsequently showed total breeding failure versus a successful breeding attempt. Similarly, physiological profiles during chick-rearing did not predict which individual would or would not initiate a second clutch, be successful with that second clutch, or survive and return to breed the following year. This highlights a marked contradiction with the widely held views that costs of reproduction are presumably associated with higher levels of reproductive investment, and are widely assumed to involve physiological costs (21). Alternatively, if birds can adjust multiple individual components of their physiology independently, there might be many physiological paths to fitness. Direct links between physiology, performance and fitness may exist, but they might be very difficult to detect without carefully designed experiments and/or more complex analytical approaches (but see companion paper, Cohen et al submitted). In fact, recent work indicates that if birds are experimentally challenged to work harder (weights and wing-clipping), a physiological cost of reproduction is detectable (63). Thus, birds that operate in ‘normal’ circumstances may not be under the maximal motivation that elicits a physiological signal linked to a cost of reproduction.

In conclusion, we were unable to detect robust relationships between physiological variables across a variety of systems and parental workload or fitness costs (immediate or future). Thus, marked individual variation in life-history traits was independent of large-scale variation in physiological phenotype at the univariate level. Our explanation of the lack of relationship between physiology and costs of reproduction revolves around several concepts. Selection on physiological traits may be filtered through behavior. Individuals may be able to independently regulate different components of their physiology, thus compensating for differences in performance and making relationships difficult to detect. Additionally, individuals experience variable ecological circumstances and may not be exerting maximal effort, further inhibiting the detection of clear relationships between physiology, behavior and current and future reproductive success in a free-ranging animal.

## Author contributions

TW and AC: conceptualization. TW and MF: investigation. TW, MF, and AC: data curation. MF and TW: formal analysis. AC and MP validation. TW: project administration and supervision. TW and AC: funding acquisition. TW, MF, AC and MP: writing-original and writing-review.

## Acknowledgements

Many thanks Allison Cornell for assistance in the field and lab. Additionally, thanks to Sarah Gray and James Hou for help with wing measurements and reticulocyte counts. Kevin Matson, Chris Harris, Oliver Love, David Costantini, Sarah Guindre-Parker, Christopher Guglielmo and David Swanson all provided invaluable advice regarding troubleshooting assays.

## Supporting Information

**Supplemental Material S1.** Physiological assay methods, quality control and data transformations.

**Supplemental Table S1.** Raw data values for body mass and physiological traits for female European starlings sampled during incubation and during chick-rearing for first and second broods in 2013 and 2014.

**Supplemental Table S2.** Correlation matrix for incubation physiology values.

**Supplemental Table S3.** Correlation matrix for chick-rearing physiology values.

